# Implementing solid-phase-enhanced sample preparation for Co-Immunoprecipitation with Mass Spectrometry for *C. elegans*

**DOI:** 10.1101/2021.08.10.455789

**Authors:** Gülkiz Baytek, Oliver Popp, Philipp Mertins, Baris Tursun

**Author notes:** Corresponding authors: Baris Tursun (BT), Gülkiz Baytek (GB).

## Abstract

Studying protein-protein interactions *in vivo* can reveal key molecular mechanisms of biological processes. Co-Immunoprecipitation followed by Mass Spectrometry (CoIP-MS) allows detection of protein-protein interactions in high-throughput. The nematode *Caenorhabditis elegans (C. elegans)* is a powerful genetic model organism for *in vivo* studies. Yet, its rigid cuticle and complex tissues require optimization for protein biochemistry applications to ensure robustness and reproducibility of experimental outcomes. Therefore, we optimized CoIP-MS application to *C. elegans* protein lysates by combining a native CoIP procedure with an efficient sample preparation method called single-pot, solid-phase-enhanced, sample preparation method (SP3). Our results based on the subunits of the conserved chromatin remodeler FACT demonstrate that our SP3-integrated CoIP-MS procedure for *C. elegans* samples is highly accurate and robust. Moreover, in a previous study (Baytek et al. 2021), we extended our technique to studying the chromodomain factor MRG-1 (MRG15 in human), which resulted in unprecedented findings.

**Method Summary:** Combination of cryo-fracture with single-pot, solid-phase-enhanced, sample preparation (SP3) to perform Co-Immuno-Precipitation followed by Mass Spectrometry (CoIP-MS) provides robust assessments of protein-protein interaction using *C. elegans* whole animals.

## Introduction

In living organisms, proteins are essential components of cellular structures, transport machineries, and perform vital enzymatic reactions during biochemical processes. Furthermore, proteins have central functions for gene expression and DNA maintenance. Therefore, studying protein-protein interactions is important to understand the vast array of molecular mechanisms and biochemical pathways in living cells.

While *in vitro* applications such as protein pull-downs indicate potential interactions, the detection of protein-protein interactions directly from cells is critical to obtain relevant insight into actual protein interaction networks. The application of Co-Immunoprecipitation (CoIP) followed by Mass Spectrometry (CoIP-MS) allows detection of in vivo protein-protein interactions. However, CoIPs from multicellular organisms are not straightforward due to various reasons depending on the research model.

The nematode *Caenorhabditis elegans* (*C. elegans*) is a powerful genetic model organism. Still, its rigid cuticle and complex tissues require optimization for protein biochemistry applications to ensure reproducibility of experimental outcomes. Therefore, we optimized CoIP-MS application to *C. elegans* by combining a native CoIP procedure with an efficient sample preparation technique called ‘single-pot, solid-phase-enhanced, sample preparation method (SP3)’ (Hughes et al. 2014, 2019).

Standard native Co-IP protocols for *C. elegans* involve physical and chemical shearing to break up the tough cuticle layer. For physical shearing, an instant freeze step in liquid nitrogen preserves protein interactions during the following cryo-fracture of the animals. Cryo-fracture makes tissues accessible for buffers containing required chemicals such as detergents to solubilize proteins prior to Mass Spectrometry (MS) analysis (Fonslow 2014). However, detergents and other components of lysis buffers that need to be used for the lysis of rigid *C. elegans* tissues strongly interfere with MS analysis resulting in reduced reproducibility of experiments (Sielaff et al. 2017). After proteins are released from the tissues, a sonication step is necessary for the fragmentation of viscous DNA to prevent interference with the target protein’s precipitation. Removing excessive DNA is especially important when purifying chromatin-regulating proteins because unspecific interactions could be mediated via genomic DNA. To distinguish specific interactions from unspecific binding proteins, which can cause significant background and noise, the immune-precipitated (IP) samples are compared with a proper negative control (Vermeulen, Hubner, and Mann 2008). Nonspecific background contaminants can be caused either by the affinity of unspecific proteins for the solid matrices used to precipitate the target protein, or due to cross-reactivities of antibodies. Having unique and efficient antibodies for the target protein of choice is not always feasible, which can be bypassed by fusing epitope tags such as the HA or FLAG to the target protein (Brizzard 2008). Some commercially available antibodies against such epitopes provide high-affinity binding allowing stringent washing procedures during the purifications steps to remove background binders. Additionally, magnetic beads that are already coupled to, e.g., anti-HA antibodies or Protein A/G with high affinities for primary antibodies (immunoglobulins) allow magnetic separation of the target protein and its interacting proteins in a highly efficient and specific manner. Again, strong detergents and denaturing conditions need to be used to elute the target protein from the beads. However, as mentioned earlier, detergents are problematic as they are incompatible with the enzymes used for proteolysis in bottom-up proteomics and MS analysis. Therefore, various methods, including ultrafiltration (Wiśniewski et al. 2009) and precipitation (Wessel & Flügge, 1984), have been utilized to purify the protein samples prior to MS analysis. Yet, these methods have shortcomings as they usually require relatively high amounts of sample and are therefore not suited for high-throughput sample preparations. To address this, a technique called *single-pot, solid-phase-enhanced, sample preparation method* = SP3 was developed as a rapid and efficient way of protein purification compatible with a range of chemicals without being restricted to high-input material (Hughes et al. 2014; Moggridge et al. 2018). SP3 makes use of carboxylated magnetic beads with a hydrophilic surface to confine proteins and peptides similarly to hydrophilic interaction liquid chromatography (HILIC) (Alpert 1990). In addition, aggregation of insoluble proteins on carboxylated beads under high organic solvent conditions has been demonstrated as a binding mechanism (Batth et al. 2019). The proteins trapped on the beads can then be washed vigorously to eliminate contaminants, detergents, and salts that interfere with MS.

To improve CoIP-MS analysis of protein-protein interactions in whole animal lysates of *C. elegans*, which contain a number of strong detergents and salts, we established a CoIP-MS working pipeline with integrated SP3 application. Additionally, we tested the effects of enzymes that remove DNA and RNA, which can cause background. We demonstrate based on the subunits of the heteromeric chromatin remodeler FACT (FAcilitates Chromatin Transcription) (Orphanides et al. 1998, 1999; Kolundzic et al. 2018) that our SP3-integrated CoIP-MS procedure for *C. elegans* samples is highly accurate and robust. In a previous study (Baytek et al. 2021), we extended our technique to studying the chromodomain factor MRG-1 (MRG15 in human), where we found that MRG-1 is SUMOylated.

## Material and Methods

### Worm strains

The wild-type *C. elegans* Bristol strain (N2) and CRISPR strains were maintained according to the standard protocol (Stiernagle 2006) at 20°. BAT1753 *hmg-3 (bar24[hmg-3::3xHA]) I, (CRISPR/Cas9)*, BAT1954 *hmg-4(bar30[hmg-4::3xHA]) III (CRISPR/Cas9)*, wild type *N2.*

### Synchronized worm population

Synchronized worms were obtained by standard bleaching procedure using sodium hypochlorite solution to disintegrate gravid adult worms as previously described (Ahringer 2006). Briefly, 5% sodium hypochlorite solution was mixed with 1 M NaOH and water in the 3:2:5 ratio. M9 buffer was applied to wash off the worms from NGM plates. Worms in M9 buffer were mixed with bleaching solution for 5 min in a 1:1 ratio, and vortexed till the adults started dissolving. To remove bleach solution completely, released embryos were washed three times with M9 buffer. After an overnight incubation, synchronized L1-staged worms were obtained. L1s were applied directly onto regular NGM plates for further maintenance of synchronized population.

### Western blot

Input and Co-IP samples were frozen at −20°C. Right before loading, SDS/PAGE sample buffer was added. Samples were boiled for 10 min to denature the proteins and centrifuged for 10 min at full speed. HMG-3::3XHA and HMG-4::3xHA were detected with anti-HMG-3/-4 antibody (Pineda) at a dilution of 1:1000.

### Antibodies and affinity matrix

anti-HMG-3/-4 antibody (Pineda); ChIP Grade anti-HA antibodies (Abcam ab9110); Sera-Mag A beads (Thermo Scientific; CAT No. 0;9-981-121, Magnetic Carboxylate Modified); Sera-Mag B beads (Thermo Scientific; CAT No. 09-981-123, Magnetic Carboxylate Modified);

### IP-MS

For each condition three biological replicates were collected as 300 μL of L4/YA staged worm pellet. Wild-type, HMG-3::3xHA, and HMG-4::3xHA worms were collected in M9 buffer, washed four times with M9 to remove bacteria, and concentrated into worm pellet after the last wash. The worm pellet was added dropwise into liquid nitrogen with special attention that the resulting “worm beads” were around the size of black pepper to ensure even grinding afterward. With the help of a pulverizer the frozen worms were then cryo-fractured. To achieve even grinding of all the tissues, worms were further ground using a mortar and pestle on dry ice. The worm powder was mixed with 1.5× of lysis buffer (20 mM HEPES pH 7.4, 150 mM NaCl, 2 mM MgCl2, 0.1% Tween 20, and protease inhibitors), dounced with tight douncer 30 times, and sonicated using a Biorupter (six times 30 sec ON, 30 sec OFF; high settings). The resulting worm lysis was centrifugated at 16,000 × g at 4° for 10 min to remove the insoluble pellet. The supernatant was transferred to 2 ml Eppendorf tubes. The protein concentration of each worm lysis for each biological replicate was determined by Bradford assay and set to 2mg/ml. Then ChIP Grade anti-HA antibodies (Abcam ab9110) was added to the samples to incubate for 30 min on a rotator at 4°C. Next, μMACS ProteinA beads (Wright 1989) (Miltenyi Biotec) were added into samples as instructed in the kit, and samples were incubated for 30 min at 4° rotating. Meanwhile, the μMACS columns were placed to magnetic separator to be equilibrated and ready for sample application. Samples were diluted 5x of their volume with lysis buffer adding up to 10 ml before being applied to columns, and the columns with bound proteins were washed three times with lysis buffer to remove background binders. The proteins were eluted with elution buffer (100 mM Tris-Cl pH 6.8, 4% SDS, 20 mM DTT), heated to 95°. Eluted samples were prepared for mass spectrometry measurements by SP3 (Hughes et al. 2014, 2019). After the final elution step, the protein amount of CoIP samples in SDS buffer was determined by DC assay that is compatible with SDS to a final amount of 50μg/μL before SP3 cleanup.

### Solid-phase-enhanced, sample preparation method = SP3

Sera-Mag A beads (Thermo Scientific; CAT No. 09-981-121, Magnetic Carboxylate Modified) and Sera-Mag B beads (Thermo Scientific; CAT No. 09-981-123, Magnetic Carboxylate Modified) were brought to room temperature for 10 min. A volume of 20 μL Sera-Mag A beads were combined with 20 μL of Sera-Mag B beads and washed with 160 μL of water by placing the water-bead mixture on a magnetic rack for PCR tubes (DynaMag PCR Magnet, Thermo Scientific, Cat: 492025), and beads were settled for 2 minutes. Magnetic beads were rinsed with 200 μL of LC-MS grade water (Thermo Fisher) by pipette mixing (off the magnetic stand). That was repeated two additional times. The final bead pellet was stored in 100 μL of water in the fridge until used. An amount of 50 μg of Co-IP sample was transferred to a PCR tube and incubated with 1 μL of benzonase (Sigma, cat. no. E8263) at 37°C for 30 min to shear and digest DNA. A volume of 10 μL of 50 mM TCEP in 50 mM ABC was added for reduction and incubated at 25°C for 20 min. 10 μL of 400 mM CAA in 50 mM ABC was added for the alkylation of the samples and incubated at 25°C for 30 min in the dark. 5 μL of the bead stock was added to each sample. For buffer exchange, acetonitrile was added to a final percentage of 50% (v/v) and incubated for 10 min off the magnetic rack while vortexing with “Vortex Genie” (manufacturer) at level 4 with 1 min intervals by avoiding spillovers from one tube to another. Then the samples were Incubated on a magnetic rack for 2 min., and the supernatant was discarded. Then 80% (v/v) 200 μL ethanol was added, and the samples were incubated off the magnetic rack for 30 sec. The samples were then incubated on a magnetic rack for 2 min, and the supernatant was discarded. Another round of washing step was carried out. 180 μL acetonitrile was added and incubated for 15 sec off the magnetic rack, followed by 2 min incubation on a magnetic rack. The supernatant was discarded, and the samples were air-dried in a safety hood. The beads were reconstituted in 5 μL digestion solution: 50 mM HEPES plus Trypsin/LysC mix (1:25 enzyme to substrate ratio, 1 μg Trypsin and 1 μg LysC per sample) and incubated for 14-16 hours at 37°C (PCR-machine with 80°C heated lid). For peptide recovery, tubes were placed on the magnetic rack for 2 min, and the supernatant was collected into fresh tubes. 40 μL 50 mM HEPES was added to the beads, resuspended, sonicated for 3 min, and placed on a magnetic rack. The supernatant was collected and combined with the first supernatant. The sample was acidified with trifluoroacetic acid (final concentration 1% (v/v)). To remove any salts and contamination stage tips with two layers of C18 were prepared as described before (Rappsilber, Ishihama, and Mann 2003). To condition the stage tips 100 μL MEOH; 100μL 50% (v/v) acetonitrile 0.1% (v/v) formic acid and 100 μL 0.1% (v/v) formic acid were used, respectively. Then the samples were loaded to the stage tips followed by washes with 100 μL 2% ACN 1%TFA and 100μL 1% FA. The stage tips were either stored at 4°C till used or eluted immediately with 50% Acetonitrile 0.1%FA. The peptides were concentrated and dried by using a SpeedVac at 35°C for app. 30 min. Concentrated peptides were resuspended in 5 μL (0.1% TFA) and a volume of 2 μL was used for injection into liquid-chromatography coupled mass spectrometry.

### LC-MS/MS analysis

Samples were measured with 1 h gradient at 15k resolution and 100ms injection time by LC-MS/MS on a Q Exactive Plus mass spectrometer (Thermo) connected to an EASY-nLC system (Thermo). The samples were separated on a 44 min gradient ramping from 5 to 55% acetonitrile using an in-house prepared nano-LC column (0.074 mm x 250 mm, 3 μm Reprosil C18, Dr Maisch GmbH) and a flow rate of 250 nl/min. MS acquisition was operated at an MS1 resolution of 70,000 and a scan range from 300 to 1700 m/z. For data-dependent MS2 acquisition the top 10 peaks were selected for MS2 with a resolution of 17,500, a maximum injection time of 60 ms and an isolation window of 2 m/z. AGC target was set to 2.5e3 and dynamic exclusion was specified to 20 sec.

### Mass Spectrometry data analysis

Raw data files were processed with default settings (unless stated otherwise) in MaxQuant version 1.5.2.8. (Cox and Mann 2008). Protein quantification was performed using the label-free quantification (MaxLFQ) algorithm (Cox et al. 2014). The “match between runs” option was chosen for transferring MS/MS identifications between LC-MS/MS runs. Enzyme Trypsin/P was used in the specific enzyme setting. Cysteine carbamidomethylation was used as fixed modification; oxidation of methionine and acetylation of the protein N terminus were set as variable modifications. Minimal peptide length of amino acids was set to 7 and a maximum of two missed cleavages were allowed. The resulting “proteinGroups.txt” was then processed using an online tool (Singh, Hein, and Stewart 2016) (Singh, Hein, and Stewart 2016).

## Results

### General method overview

Interaction proteomics involves detecting the specific interactors of a particular protein when those interactors are significantly enriched. For this purpose, label-free purification methods provide sufficient robustness with the implementation of the label-free quantification (LFQ) algorithm in MaxQuant software (Cox et al. 2014). When label-free approaches, such as MaxLFQ, were compared to metabolic labeling, MaxLFQ proved to be as accurate as, for example, SILAC (Eberl et al. 2013). Overall, LFQ analysis is suitable for interaction proteomics, but accurate assessment of individual protein ratios in LFQ requires a t-test with three or more replicates (Cox et al. 2014). Therefore, we performed at least three biological replicates for each bait protein. During method optimization, manual cryofracturing was performed using a mortar and pestle (Figure 1).

**Figure 1:**
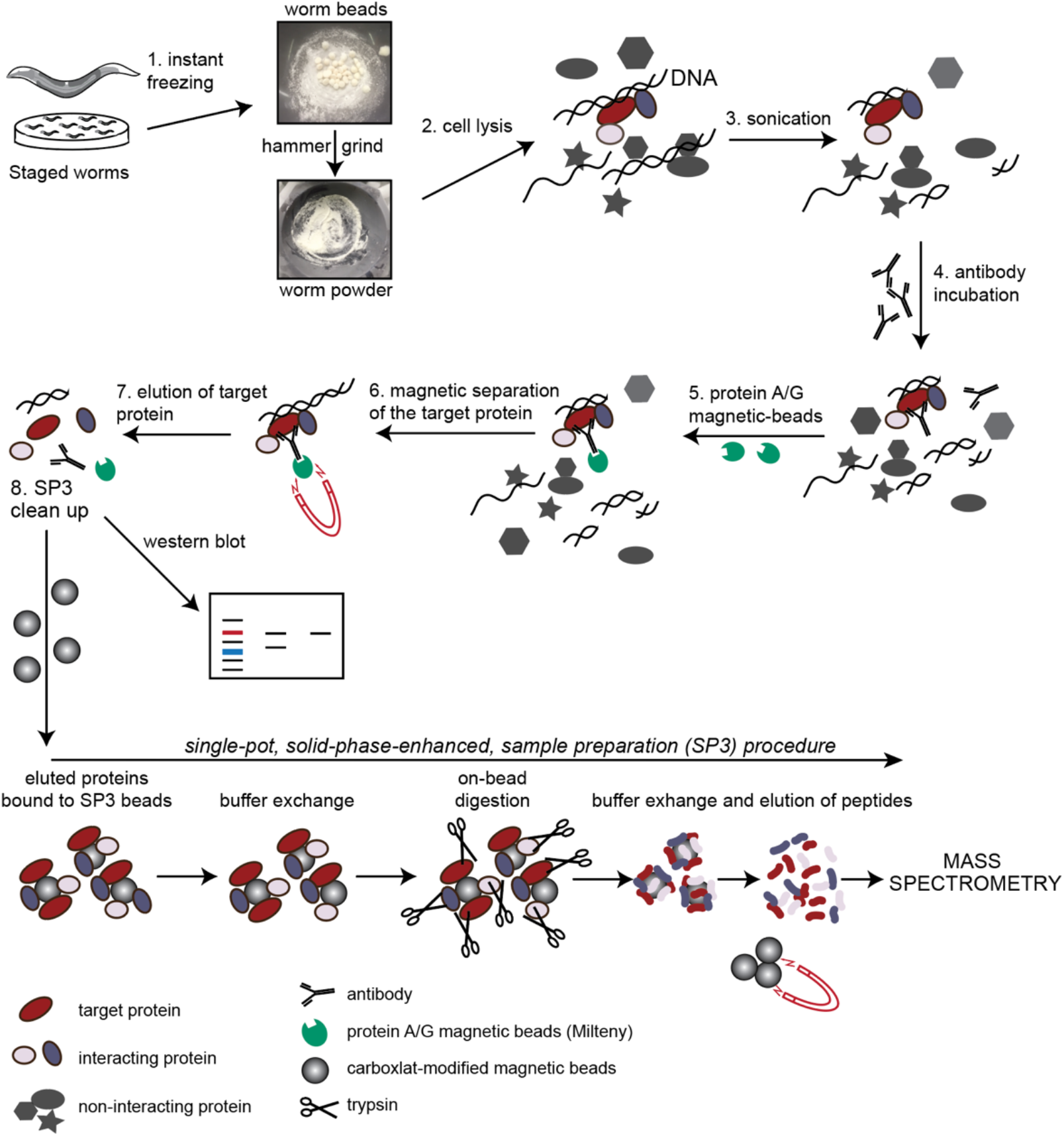
Workflow of native CoIP-MS for *C.elegans* lysates with SP3. See main text and Material and methods for details.

For cryofracturing worms were first frozen in liquid nitrogen as pellets and subsequently powdered using hammering and grinding followed by cell lysis and sonication (**Figure 1**, see Material and Methods for details). For our endogenously expressed 3xHA-tagged proteins (derived by CRISPR/Cas9 gene editing), samples were incubated with anti-HA antibodies followed by protein A/G incubation, which are coupled to magnetic beads and therefore allow efficient separation of the beads with bound protein using magnetic racks (**Figure 1**, see Material and Methods for details). After eluting the proteins off the immunoprecipitation beads using an SDS and DTT-containing elution buffer, the protein fraction was further cleaned up using the solid-phase-enhanced, sample preparation method (SP3) (Hughes et al. 2014, 2019). SP3 makes use of carboxylated magnetic beads with a hydrophilic surface to bind proteins and peptides, thereby allowing vigorous washing steps to eliminate contaminants, detergents, and salts that interfere with MS. Another aspect of removing noise-causing cell components prior to performing the immunoprecipitation is removing DNA and RNA. Nucleic acids can cause artificial interaction of DNA or RNA-binding proteins by bridging two proteins that both bind nucleic acids but do not interact directly with each other (Fiil et al. 2008). DNA and RNA can be eliminated from cell lysates by applying enzymes such as DNase or benzonase, which cleaves all DNA and RNA (Fiil et al. 2008). However, it is not clear whether their use can interfere with CoIP-MS. Therefore, we assessed, as described in the next section, their effect on CoIP-MS. We used whole worm samples and targeted the chromatin-binding protein HMG-3, which was previously tagged with 3xHA using CRISPR/Cas9 (Kolundzic et al. 2018).

### The effect of benzonase and DNase on the interactors of chromatin regulator HMG-3

Co-IPs were performed with or without DNase and benzonase treatment, separately. Benzonase was chosen for being a more efficient enzyme compared to DNase and its ability to chop down both DNA and RNA molecules at 4°C. In order to check the reaction efficiencies of both enzymes at 4°C, which is the temperature that the IPs are carried out, 5μg genomic DNA (gDNA) was incubated with each enzyme (at optimal conditions according to manufacturer) and compared with a longer reaction period at 4°C **(Figure 2A.** Benzonase showed a higher efficiency by digesting the whole gDNA at 4°C. Although there was gDNA detectable after DNase application, less DNA after 16h at 4°C compared to DNase’s reported optimal reaction condition was an indication of the enzyme’s activity at 4°C **(Figure 2A).** In order to compare the impact of benzonase and DNase treatment on chromatin regulator interactions, protein-protein interactions of the germline-specific chromatin regulator HMG-3 was compared upon enzyme treatments and the control. In the control case, 41 proteins appeared to be interacting with HMG-3 significantly **(Figure 2B)**. Gene ontology analysis based on the wormbase enrichment analysis (https://wormbase.org/tools/enrichment) revealed unexpected enrichments such as for actin-binding or actin-filament-based processes **(Figure 2C).** This might be due to HMG-3’s unspecific interactions with structural proteins due to the presence of the DNA. Upon DNAse treatment **(Figure 2D)**, gene ontology (GO) term analysis of the significant interactors of HMG-3 showed the expected involvement in more chromatin-based and germline-specific functions such as reproduction **(Figure 2E).** The elimination of DNA-directed unspecific interactions of HMG-3 upon DNAse treatment could account for a reduction of background interactions. Also, benzonase-applied samples showed a different enrichment of protein interactions **(Figure 2F)**. Even though the analyzed protein amounts were the same in all the samples, the number of interaction partners and overall identified proteins dropped significantly for benzonase treated samples **(Figure 2F)**. This might be due to either the higher potency of benzonase or the degradation of RNAs in addition to DNA upon benzonase application **(Figure 2G).** Overall, the milder condition of DNase treatment to preserve potentially relevant interactions was found to be more suitable for pull-down assays of chromatin-binding proteins.

**Figure 2:**
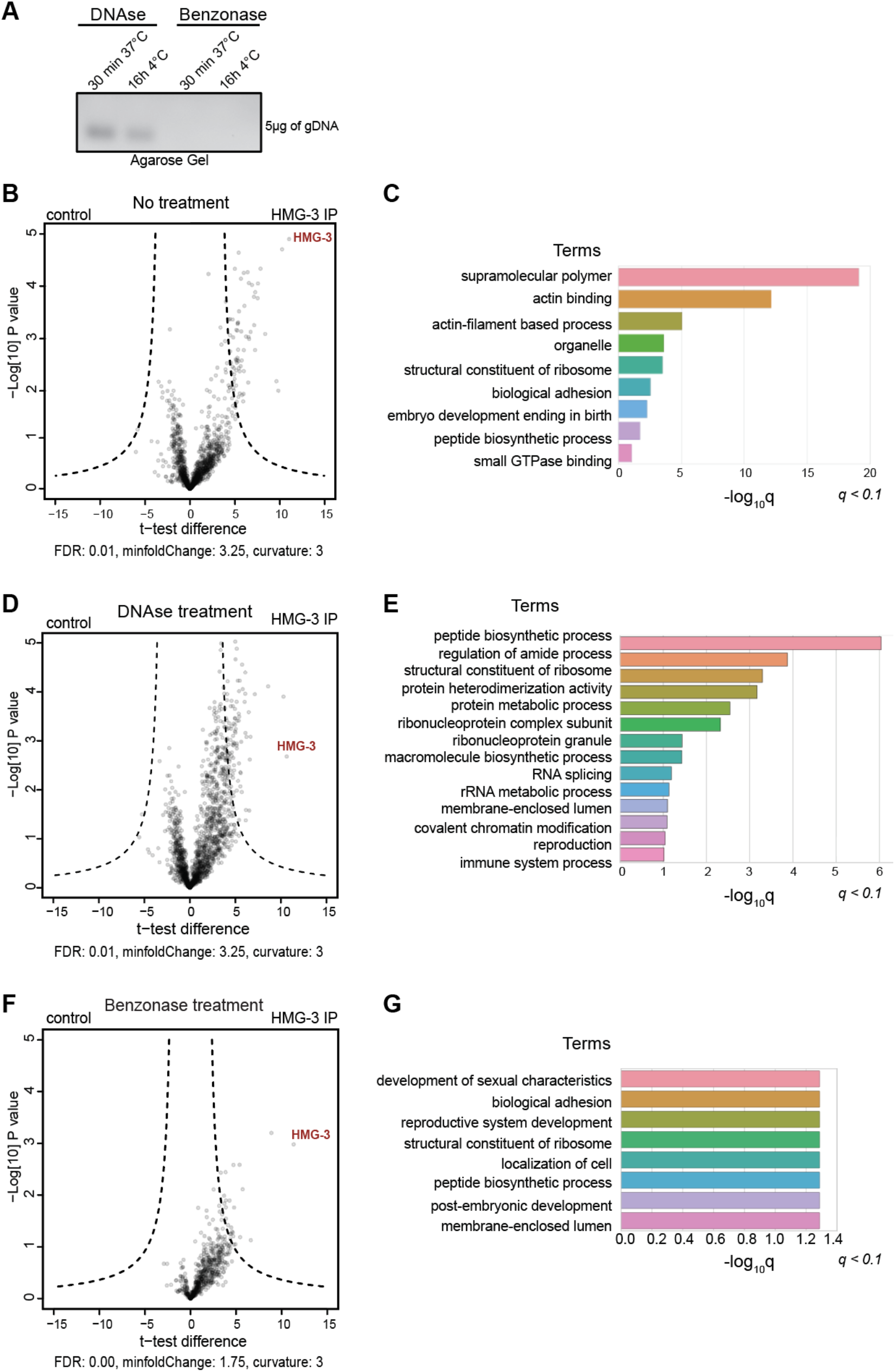
The impact of DNase and benzonase on detectable interactions of the chromatin regulator HMG-3: Co-immunoprecipitations (co-IPs) with subsequent mass spectrometry (IP-MS) to assess HMG-3’s protein interactions. **A)** Agarose gel-electrophoresis detection of 5μg genomic DNA (gDNA) incubated with DNase and benzonase, 37°C and 4°C for 30 min and 16h, respectively. Wild type (N2) was used with anti-HMG-3 and unspecific antibodies as control. **(B, D, and F).** Volcano plots showing statistically significant enrichment of co-precipitated proteins. Statistics: t-test, adjusted P-value is set as indicated on each plot as false discovery rate cut-off. **(B, E, and G)** Gene set enrichment analysis using wormbase version WS279 of proteins interacting with HMG-3.

### Testing the specificity and cross-reactivity of anti-HA antibody for CoIP-MS

In several affinity purification studies, large epitope tags were shown to be more likely to affect the function of the proteins by changing their conformation and folding (Kimple, Brill, and Pasker 2013). Moreover, CRISPR knock-in of fluorescent proteins is known to be less efficient than smaller tags such as HA (Dokshin et al. 2018).

Aiming to assess the efficiency of the small affinity purification based on the HA tag, we made use of strains tagged with 3xHA-tag at the C-terminus of the FACT complex members HMG-3 and HMG-4 **(Figure 3)**. FACT is a chromatin regulator which was identified as a reprogramming barrier both in *C. elegans* and human (Kolundzic et al., 2018). FACT’s interaction partners were previously not analyzed in *C. elegans*. FACT, <u>fa</u>cilitates <u>c</u>hromatin transcription, which is composed of SSRP1/SUPT16H in human (Orphanides et al. 1998, 1999; Kolundzic et al. 2018), forms two different complexes based on the tissue type in *C. elegans* (Kolundzic et al., 2018). FACT is composed of HMG-3 and SPT-16 in the germline with a minor presence of HMG-4 in the germline as well, while in somatic tissues HMG-3 is completely absent and substituted by HMG-4 (Kolundzic et al., 2018). Keeping in mind the tissue specificity of the heterodimers of FACT in *C. elegans* (Kolundzic et al., 2018) immunoprecipitating HMG-3 and HMG-4 with the same antibody to compare their interaction partners is an ideal set up to evaluate the efficiency of our CoIP-MS protocol.

**Figure 3:**
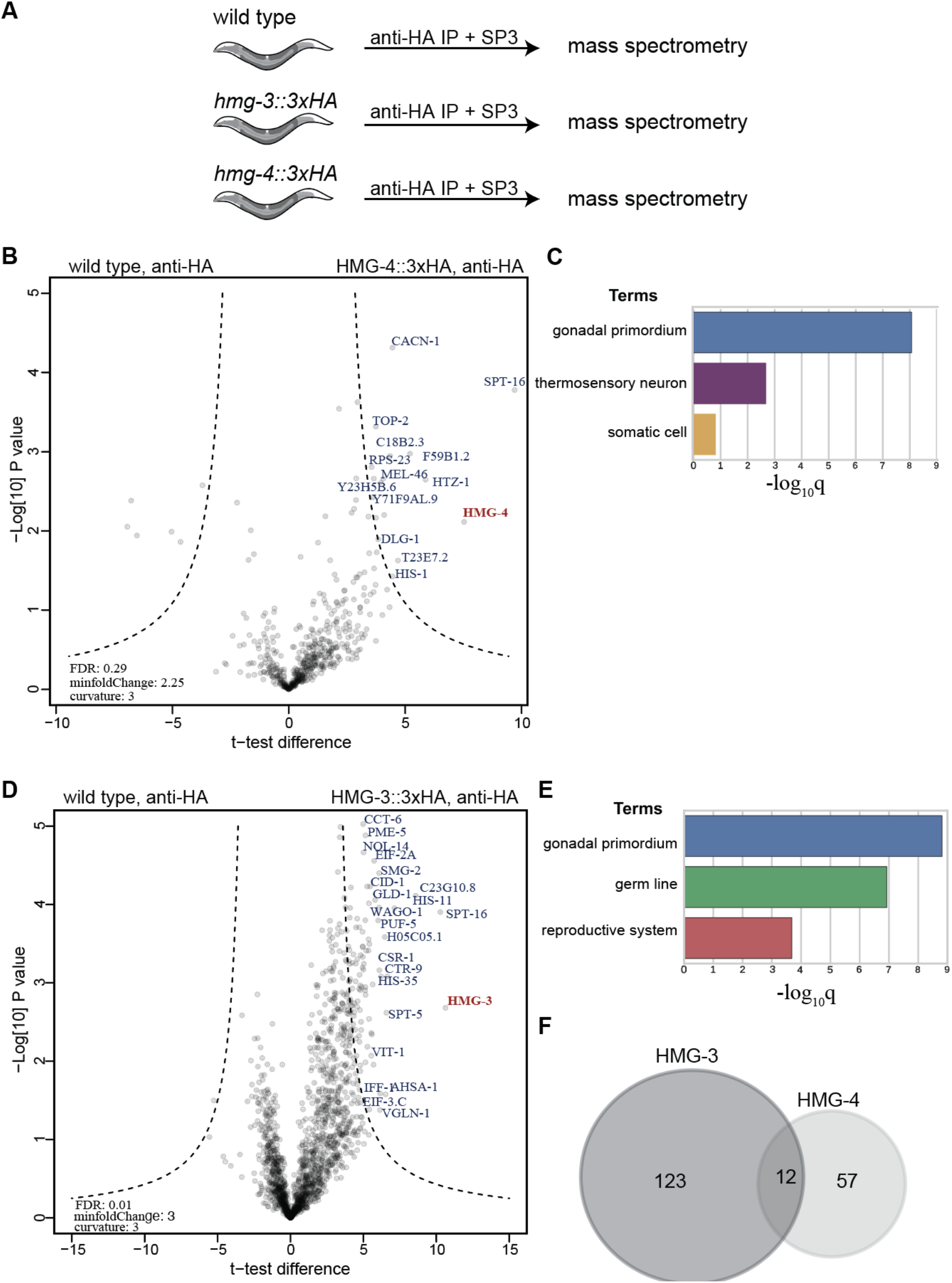
Revealing tissue-specific FACT-interacting proteins in *C. elegans*: **(A)** The protein extracts of both the wild type N2, HMG-3::3xHA, and HMG-4::3xHA strains were incubated with ChIP grade anti-HA antibody. **(B and D)** Volcano plots showing specific protein interactions of HMG-3::HA and HMG-4::3xHA based on pull-down experiments of the three biological replicates, respectively. HMG-3 and HMG-4 interaction partners are written in blue, HMG-3 and HMG-4 shown in red. The stringency cut-offs (hyperbolic curves) are drawn with a 0.01 false discovery rate as indicated on the volcano plot. **(C and E)** Tissue enrichment analysis of proteins interacting with HMG-3 and HMG-4 using wormbase version WS279, q<0.1. **(F)** Venn diagram showing the overlapping interactors of HMG-3 and HMG-4.

To assess the robustness of the Co-IP protocol, CRISPR-edited animals carrying HMG-3::3xHA, HMG-4::3xHA, and wild-type N2 worms were prepared **(Figure 3A)**. In the pull-down assays against HMG-3::3xHA and HMG-4::3xHA, 200μL of worm pellet from young gravid hermaphrodites were collected and snap frozen in liquid nitrogen (see Material and Methods for detailed procedures). Three biological replicates were prepared in parallel for each strain. The same anti-HA antibody was used with wild-type N2 worms to detect unspecific binders of the antibody.

Co-IPs were prepared, and abundances of the co-purified proteins were assessed by label-free quantitative mass spectrometry to identify proteins specifically enriched in the HMG-3::3xHA and HMG-4::3xHA pull-downs **(Figure 3A).** The most significant interactor for both proteins was their heterodimer partner SPT-16 (Orphanides et al. 1998, 1999; Kolundzic et al. 2018), indicating the robustness of the Co-IPs **(Figure 3B–3E).** Notably, tissue ontology terms for the significantly enriched proteins for both HMG-3 and HMG-4 showed differential tissue expressions as expected. While HMG-3-enriched proteins were correlated exclusively with the germline, HMG-4 interacting proteins were correlated with gonadal primordium and somatic cells such as neurons **(Figure 3C, and 3E)**. 12 proteins were identified to interact with both HMG-3 and HMG-4 **(Figure 3F).** Overall, these results revealed previously unknown potential interaction partners of FACT, demonstrating the high efficiency of our CoIP-MS method by maintaining the tissue-specific interaction partners of HMG-3 and HMG-4.

## Conclusion

Physiological processes in living organisms depend on protein interaction. Hence, studying the *in vivo* interactome of proteins is required to expand our knowledge on protein dynamics derived from investigating cell lines outside of their physiological environments (Rual et al. 2005; Papachristou et al. 2018). To establish a reliable method for studying protein interactions in a multicellular organism, we used the nematode *C. elegans*. While mass spectrometry for protein interaction studies was conducted in *C. elegans* previously (Fonslow 2014; Moresco 2012; Chen et al. 2016), its application to worms is not straightforward. Proper lysis of all tissues is difficult due to the cuticle, and the content of the whole worm body, such as fat and other organic compounds, can severely confound reproducibility.

In this report, we are providing a highly reproducible and robust protocol for CoIP-MS. We combined a native Co-IP protocol for worm lysates with a single-pot, solid-phase-enhanced sample preparation method termed SP3 (Hughes et al. 2014, 2019; Moggridge et al. 2018). We introduced the 3xHA tag, which can be targeted using commercially available anti-HA antibodies in a highly efficient manner, by CRISPR to two proteins of interest, HMG-3, and HMG-4. In a parallel study, which is reported in an accompanying manuscript, we also tagged the chromatin regulator MRG-1 with 3xHA using CRISPR editing (Baytek et al. 2021). This strategy allows that the wild-type animals without the knock-in can be used to identify the background binders of the antibody and purification resin. In order to have confidence in the data, three replicate experiments were performed for HMG-3 and HMG-4, while in the accompanying study, there were at least 10 replicates for MRG-1. We also evaluated the effects of benzonase and DNase treatment of the protein lysate sample preparation. Both enzymes can be used to remove nucleic acids as these can cause background protein interactions of, for instance, chromatin-binding proteins such as HMG-3 and HMG-4. Interestingly, the number of protein interactions and intensities of detected peptides for HMG-3 were significantly less for benzonase-treated samples. While it remains to be understood whether this effect is due to the activity of benzonase also on RNA, we decided to apply DNase for further analysis, since many functionally relevant interactions of chromatin-binding proteins also depend on the presence of RNA and DNA. DNAse treatment may preserve potentially relevant interactions that may become otherwise undetectable due to decreased peptide intensities upon benzonase treatment.

To measure the intensities of the precipitated proteins, we used label-free quantification. Although the detected peptides are utilized to deduce the identity and quantity of proteins in bottom-up proteomics, mass spectrometric measurements are not inherently quantitative (Aebersold and Mann 2003). In bottom-up proteomics, the mass spectrometers are usually coupled to high-performance liquid chromatography (LC). This enables the separation of complex peptide mixtures based on their molecular features before ionization and transfer into the mass spectrometer (Zhang et al. 2013). The ion signals corresponding to the peptides gathered from the mass spectrometers cannot be inferred as the absolute abundances of the protein species in a given sample for several reasons; the ionization efficiency can be immensely different for different types of peptides, the protein purification methods in use may favor specific type of proteins at the same time, causing the loss of others. Also, the instrument in use may have different ranges of efficiency on different sampling time points. To circumvent these issues, stable isotope techniques such as metabolic labeling or *in-vitro* chemical labeling may be used for relative quantification of several samples (Bantscheff et al. 2007). The power of stable isotope-based methods is to pool various samples and measure them in the same LC-MS run. Although the labeled and unlabeled peptides cannot be distinguished based on their chemical composition and chromatographic behavior, the mass spectrometry can differentiate them based on their mass difference. However, using stable isotope labeling methods is labor-intensive, time-consuming, and costly when applied in model organisms. As an alternative to label-based methods, different label-free quantification methods can be used. For instance, simple spectral counting, such as protein abundance index (PAI) (Rappsilber et al. 2002), extracted ion chromatography (XIC)-based methods by incorporating peptide ion intensities to their chromatographic profiles, adding an extra dimension to the quantification of the peptides (Grossmann et al. 2010). This feature made XIC-based methods superior to spectral counting (Grossmann et al. 2010). The advancement of label-free methods continued with the implementation of the label-free quantification (LFQ) algorithm in MaxQuant software as an accurate and robust method (Cox et al. 2014). A previous study comparing metabolic labeling and label-free approaches for interaction proteomics in a mouse cell line revealed that MaxLFQ achieved similar quantification properties to SILAC (Eberl et al. 2013), although low LFQ intensities resulted in reduced accuracy (Eberl et al. 2013). Nevertheless, for interaction proteomics, the specific interactors of a given protein are detected if their enrichment levels are significant, which makes LFQ analysis the method of choice for interaction proteomics in C. elegans in our study.

The LFQ analysis of HMG-3 and HMG-4 CoIP-MS, as well as of MRG-1 in a previous study (Baytek et al. 2021), delivered highly reproducible results. We confirmed predicted interactions such as with SPT-16 (Orphanides et al. 1998, 1999; Kolundzic et al. 2018) in a highly robust manner. Moreover, all 11 CoIP-MS for MRG-1 showed a very strong correlation based on Pearson correlations also in our previous study (Baytek et al. 2021). Overall, our protocol enables a efficient and robust examination of protein interactions in *C. elegans* based on CoIP-MS. For future applications, *C. elegans* also provides the potential to reveal tissue-specific protein interactions in particular when epitope-tagged proteins are expressed in confined tissue lineages. In addition, tissue-specific labeling techniques such as in vivo biotinylation (Waaijers et al. 2016) can be applied and interactions can be validated also via microscopy-based approaches.

## Financial disclosure and Acknowledgments

Thanks to Sergej Herzog for technical assistance and the CGC, funded by NIH P40OD010440, for providing strains. This work was partly sponsored by the ERC-StG-2014-637530 and the Max Delbrueck Center for Molecular Medicine in the Helmholtz Association. All procedures conducted in this study were approved by the Berlin State Department for Health and Social (LaGeSo). The authors declare that they have no competing interests.

